# Distribution and efficacy of ingested dsRNA targeting tissue-specific genes in the Argentine ant, *Linepithema humile*

**DOI:** 10.1101/2025.04.22.649969

**Authors:** Mathew A. Dittmann, Grzegorz Buczkowski, Brock A. Harpur

## Abstract

With the increasing availability of genomic and transcriptomic information within Formicidae, the investigation of gene function has become possible within ants. However, eusocial life history renders the generation of transgenic strains of ants difficult outside of specific ant lineages that can induce reproductive behavior in workers. RNA interference (RNAi) remains a practical option to investigate gene function within their genomic context in ant lineages that do not exhibit reproductive behavior in workers. This method can be leveraged to investigate the large odorant receptor (OR) gene families present in ant clades. However, ORs tend to be expressed in a tissue-specific fashion, and the capability of dsRNA to achieve knockdown of genes exhibiting localized expression remains uncertain. In this study, we use fluorescently labelled dsRNA to track the spread of dsRNA through the worker body, qPCR to verify that dsRNA can knockdown gene expression in the antennae, and behavioral trials to verify that knockdown of ORs affects nestmate recognition. We found that orally-administered dsRNA is capable of being spread systemically and knocking down tissue-specific genes. Additionally, we found that RNAi may be useful in investigating the influence of ORs in eusocial insect nestmate recognition.

## 2 Introduction

One of the primary advantages that allow eusocial species to function in diverse ecosystems is the ability to form colonies composed of many adults that work together for the colony’s benefit. Nestmate recognition through pheromone signaling is the backbone of colony formation and maintenance in eusocial insect species (Smith and Liebig, 2017; Sturgis and Gordon, 2011; Yan and Liebig, 2021). Workers rely on primarily hydrocarbon-based odorant cues to distinguish nestmates from non-nestmate workers (Sturgis and Gordon, 2011). These hydrocarbons are detected by odorant receptors in the antennae, embedded in the odorant receptor nerve (Suh et al., 2014). The identity of specific ORs involved in nestmate recognition is unknown, but an expansion of nine-exon ORs in Formicidae is considered a likely group (McKenzie et al., 2016; Smith et al., 2011). In particular, some of these nine-exon ORs show antennae-biased gene expression in *L. humile* (Dittmann et al., 2024). Much work has been done to investigate how cuticular hydrocarbons (CHCs) and ORs influence nestmate recognition in ant species through a variety of methods, such as environmental modification and ligand-binding assays (Ferguson et al., 2020; Liang and Silverman, 2000; Obin, 1986). However, the contribution of individual tuning ORs to nestmate recognition remains unclear, due to difficulties in studying gene function in eusocial species.

Thanks to the development of mutagenesis tools such as *CRISPR-Cas9*, functional characterization of genes has become much more approachable in recent years (Doudna and Charpentier, 2014). However, these tools only generate transgenic individuals, and rely on reproduction in the mutants to generate a population that can be used for experimentation. While some ant species are capable of worker reproduction, obligate sterility in many ant species renders this tool impractical for functional characterization (Trible et al., 2017; Yan et al., 2017). Insertion of genes into model organisms remains an option in these cases, but it the process is time-consuming and removes the gene from its organismal context (Slone et al., 2017). However, RNA interference (RNAi) is a potentially viable alternative that requires much less effort to set up experiments, and can be performed in eusocial species (Wilson and Doudna, 2013). Prior work has identified some likely candidate ORs that may be involved in nestmate recognition. In this paper, we examine the potential of RNAi in investigating the function of odorant receptors putatively involved in nestmate recognition in the Argentine ant, *Linepithema humile*.

## 3 Methods

### 3.1 Colony Collection and Maintenance

Portions of two hostile *Linepithema humile* colonies were collected from Winston-Salem and Raleigh, North Carolina via nesting substrate collection and brought to the lab. The colonies were extracted from the nesting material and allowed to move into artificial nests consisting of foil-covered test tubes. Once extracted, the colonies were transferred to Fluon-lined trays and kept under 80 ± 2°F, 50 ± 5% relative humidity, and 14:10 light:dark cycle. Colonies were provided with water, 20% sugar water, and freshly killed American cockroaches. For dsRNA feeding trials, foraging workers were collected from both colonies and transferred to Fluon-lined tubs and provided with foil-wrapped test tubes nests half-filled with water. Colony fragments were starved for 24 hours before experiments were started, at which point they were fed daily with 10% sucrose solution dosed with dsRNA.

### 3.2 dsRNA Synthesis and Administration

Worker mRNA was extracted using an SV Total RNA Isolation Kit (Z3101, Promega, Madison, WI). cDNA was generated using a SensiFAST cDNA Synthesis Kit (BIO-65054, Meridian Bioscience, Memphis, TN). DNA was extracted from *L. humile* workers using a DNEasy Blood and Tissue Kit to generate dsRNA targeting non-coding DNA (69504,Qiagen, Germantown, MD). T7 PCR products were generated using a SensiFAST SYBR No-ROX Kit (BIO-98050, Meridian Bioscience, Memphis, TN). dsRNA was generated from T7 Products using a Durascribe T7 Transcription Kit (DS010925, Biosearch Technologies, Middleton, WI). dsRNA used in histological work was treated with *Silencer* siRNA Labeling Kit with Cy3 dye (AM1632, ThermoFisher Scientific, Waltham, MA). dsRNA was prepared for trials by mixing 1:1 with autoclaved 20% sucrose solution to create a 10% sucrose working solution. The working solution was aliquoted into PCR tubes and stored at 4^°^C until needed.

### 3.3 Histology

Foragers were collected from laboratory colonies and kept in Fluon-lined plastic tubs. Colony fragments were provided artificial nests consisting of test tubes half-filled with water and wrapped in aluminum foil to provide darkness preferred by the ants. Several dozen workers were starved for 24 hours, then provided with 100 µl doses of 10% sugar water dosed with fluorescent-tagged dsRNA every other day for 6 days. Twenty-four hours after initial dosing, 5 workers were removed from the tub every day and stored in RNALater (AM7021, ThermoFisher, Waltham, MA). After 6 days, workers were taken from the RNALater and incubated for 48 hours in 10% paraformaldehyde and submitted to the Purdue Histology Core for processing and visualization. Samples were processed and embedded in paraffin, then sectioned onto charged slides and deparaffinized. The slides were stained with Alexa Fluor 555 secondary antibody and DAPI, then dehydrated and cover slips applied. The slides were then scanned at 20x for fluorescence using a Leica Aperio imaging system.

### 3.4 dsRNA Efficacy Testing

Foragers were collected from laboratory colonies and kept in Fluon-lined plastic tubs. Colony fragments were provided foil-wrapped test tubes as shelter with water sequestered behind a cotton ball. Workers were starved for 24 hours, then provided with daily aliquots of 50 µl of sugar water dosed with dsRNA (See Table 1 for concentration) for 6 days. Colony fragments from each colony were fed dsRNA that targeted either an odorant receptor gene, or a non-coding region of the *L. humile* genome, for a period of 6 days. Target genes were chosen based on prior studies showing nine-exon clade status and antennae-specific gene expression (Dittmann et al., 2024; Smith et al., 2011). After 6 days, workers were subjected to 1:1 aggression assays using a 1-4 aggression scale (Roulston et al., 2003).One worker from each colony fragment was removed from their tubs via paintbrush and transferred to a Fluon-lined 35×10mm Petri dish where they were allowed to interact. Ten interactions were scored using the aggression scale in each worker pairing, at which point each worker was transferred to a separate Fluon-lined tray for storage until aggression assays were completed. Ratings were further designated as Not Aggression (aggression levels 1 and 2) or Aggression (aggression levels 3 and 4) to generate a proportion of aggressive behavior from 0 to 1. Once aggression trials were completed, the workers were immersed in RNALater at 4^°^C for later dissection.

**Table 1.**
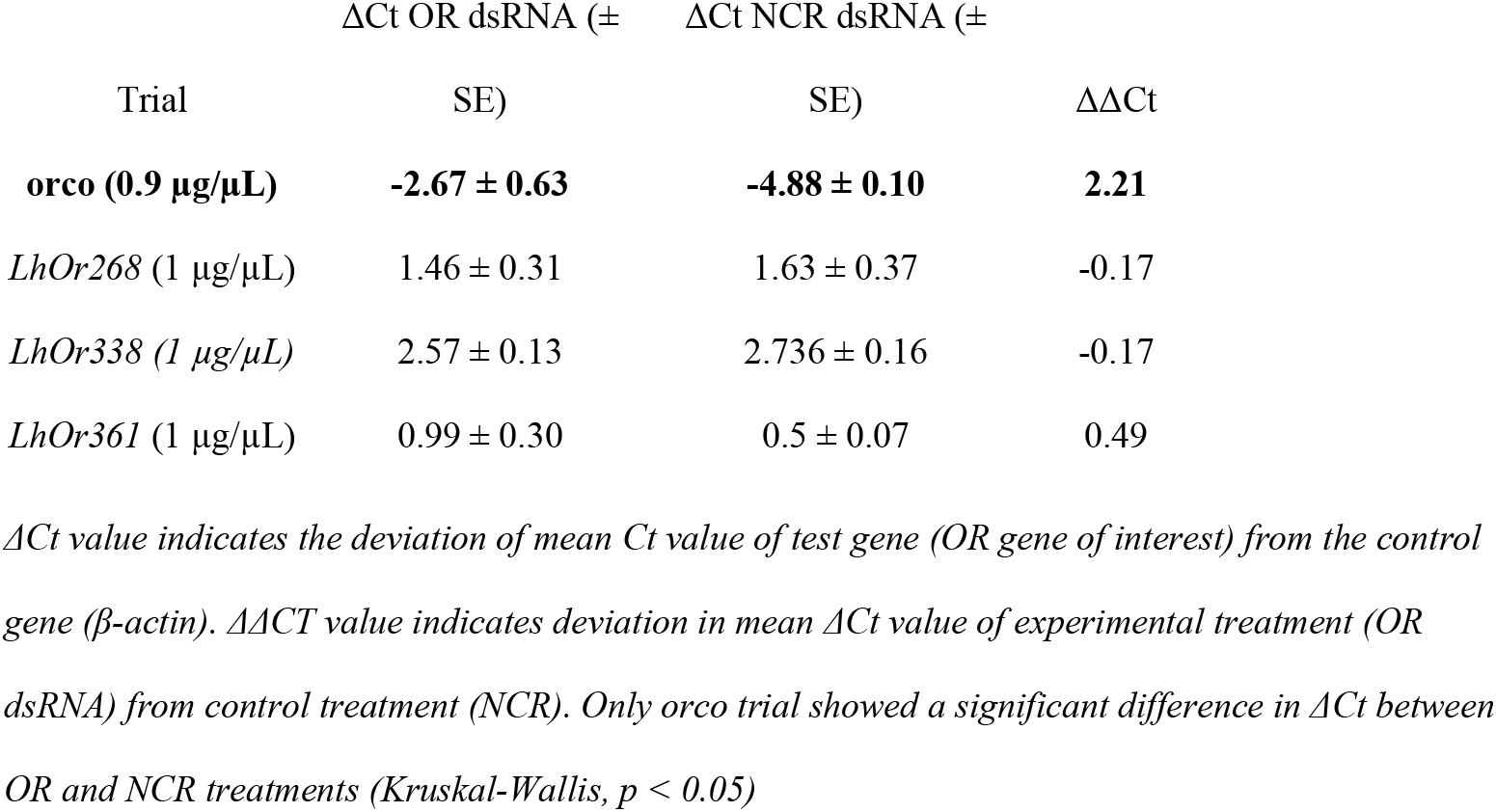
Table of qPCR data from aggression assay workers.

| Trial | $\Delta\text{Ct OR dsRNA } (\pm \text{ SE})$ | $\Delta\text{Ct NCR dsRNA } (\pm \text{ SE})$ | $\Delta\Delta\text{Ct}$ |
| --- | --- | --- | --- |
| ***orco* (0.9 $\mu\text{g}/\mu\text{L}$)** | **$-2.67 \pm 0.63$** | **$-4.88 \pm 0.10$** | **2.21** |
| *LhOr268* (1 $\mu\text{g}/\mu\text{L}$) | $1.46 \pm 0.31$ | $1.63 \pm 0.37$ | -0.17 |
| *LhOr338* (1 $\mu\text{g}/\mu\text{L}$) | $2.57 \pm 0.13$ | $2.736 \pm 0.16$ | -0.17 |
| *LhOr361* (1 $\mu\text{g}/\mu\text{L}$) | $0.99 \pm 0.30$ | $0.5 \pm 0.07$ | 0.49 |

$\Delta\text{Ct}$ value indicates the deviation of mean Ct value of test gene (OR gene of interest) from the control gene ( β-actin). $\Delta\Delta\text{CT}$ value indicates deviation in mean $\Delta\text{Ct}$ value of experimental treatment (OR dsRNA) from control treatment (NCR). Only *orco* trial showed a significant difference in $\Delta\text{Ct}$ between OR and NCR treatments (Kruskal-Wallis, $p < 0.05$ )

Both antennae were removed from workers that completed aggression trials and stored in RNALater at 4^°^C during sample collection. Forty antennae were collected from twenty workers and pooled for each sample. Once sample collection was complete, antennae were removed from RNALater and RNA was extracted using an SV Total RNA Isolation Kit, and cDNA was generated using a SensiFAST cDNA Synthesis Kit, with input RNA standardized based on input tissue. qPCR was conducted using a SensiFAST SYBR no-ROX Kit on a CFX96 Real Time System/C1000 Thermal Cycler (Bio-Rad Laboratories, Hercules, CA). (see Fig. S1 for protocol). Two technical replicates were done for each sample, and β-actin was used as the control gene. Statistical analyses of aggression assay and qPCR Ct data were conducted via Kruskal-Wallis test using *R* (4.1.0) and *RStudio* (2023.03.1 Build 446).

## 4 Results

The Cy3 signal bound to the dsRNA was strongest in the abdominal tissues, indicating that dsRNA has difficulty exiting the midgut into the hemolymph (Figure 1A). However, dsRNA can cross the gut lumen, enter the hemolymph, and reach all major body regions of worker ants within twenty-four hours of ingestion (Figure 1B). The ingestion of *orco* dsRNA resulted in significant reductions in the expression of *orco* gene in the antennae compared to noncoding dsRNA (NCR) (p < 0.05) (Table 1). However, none of the dsRNA treatments targeting tuning ORs produced a significant reduction in gene expression (Table 1). The ingestion of *orco* dsRNA also significantly reduced intercolony worker aggression (0.8 on a 0-1 aggression scale) when compared to workers fed non-coding region dsRNA (0.9 on a 0-1 aggression scale) (Figure 2A). This reduction in aggression is driven by a pattern of slight loss in aggression (0.1-0.3 reduction) in many treated ants versus control (Figure 2B). Aggression assays conducted on ants treated with dsRNA targeting *LhOr268* (0.85 vs 0.91, p = .059) and *LhOr338* (0.78 vs 0.83, p = .069) show a small but insignificant loss in aggression while *LhOr361* showed no change in aggression (.089 vs .086, p = 0.74), though none of the tuning OR trials were significant (Figure 3). Similar to the *orco* aggression trial, The reduction in aggression in *LhOr268* and *LhOr338* treatments versus their NCR controls is driven by a pattern of small reduction in aggressive behavior (0.1-0.2 reduction) across numerous 1:1 worker pairings (Figure 4).

**Figure 1.**
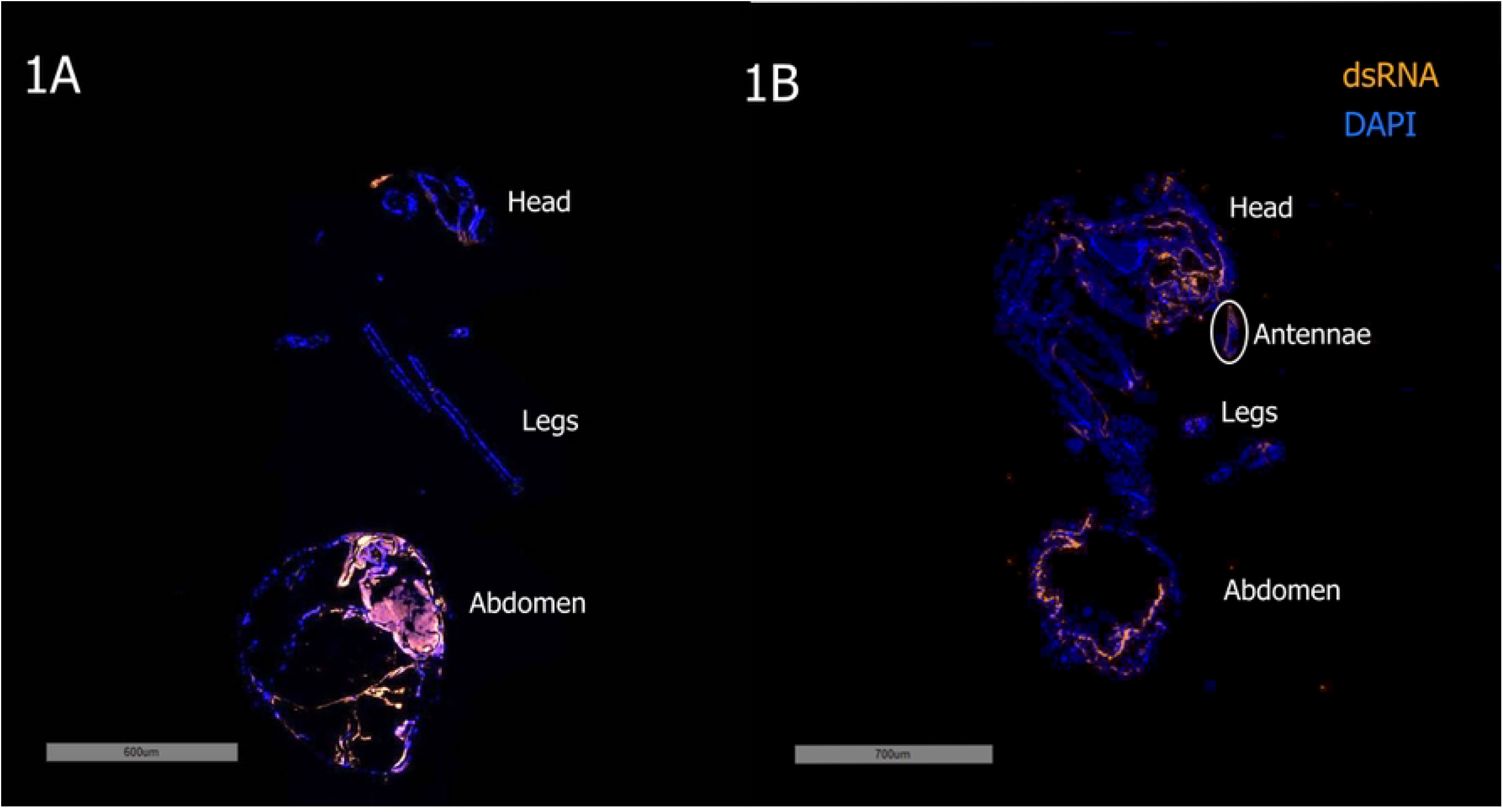
Confocal microscopy of *L. humile* longitudinal sections. (A): Day 5 longitudinal image with fluorescence showing dsRNA (Cy3-tag/orange) and cell nuclei (DAPI/blue). Very strong orange signal in abdominal tissues indicate dense concentrations of dsRNA present in worker midgut. (B): Day 1 longitudinal image with fluorescence showing dsRNA (Cy3-tag/orange) and cell nuclei (DAPI/blue). dsRNA signal is found in all major body regions of worker, indicating that dsRNA is capable of crossing midgut into hemolymph and spreading throughout body within twenty-four hours of ingestion.

**Figure 2.**
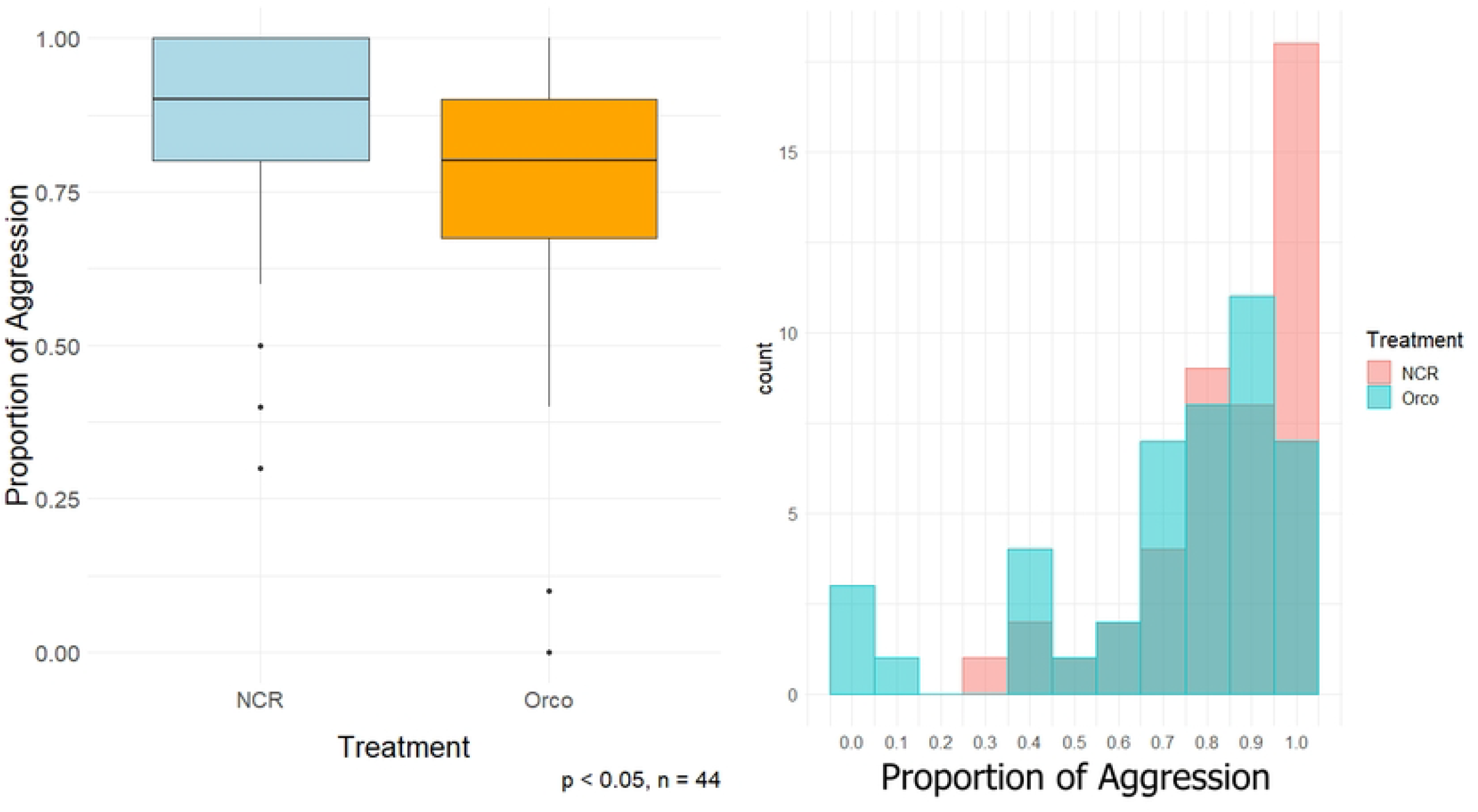
Aggression of 0.9 µg/µL *orco* dsRNA vs. NCR dsRNA. (A): Aggression assay boxplot comparing average aggression between ants treated with *orco* dsRNA vs NCR dsRNA. Ants treated with *orco* showed a significant reduction in cross-colony aggression compared to NCR-treated ants. (B): Aggression assay histogram comparing the distribution of proportion of aggression responses between ants treated with *orco* dsRNA vs NCR dsRNA. Aggression patterns shifted broadly less aggressive in ants treated with *orco* dsRNA than NCR-treated ants.

**Figure 3.**
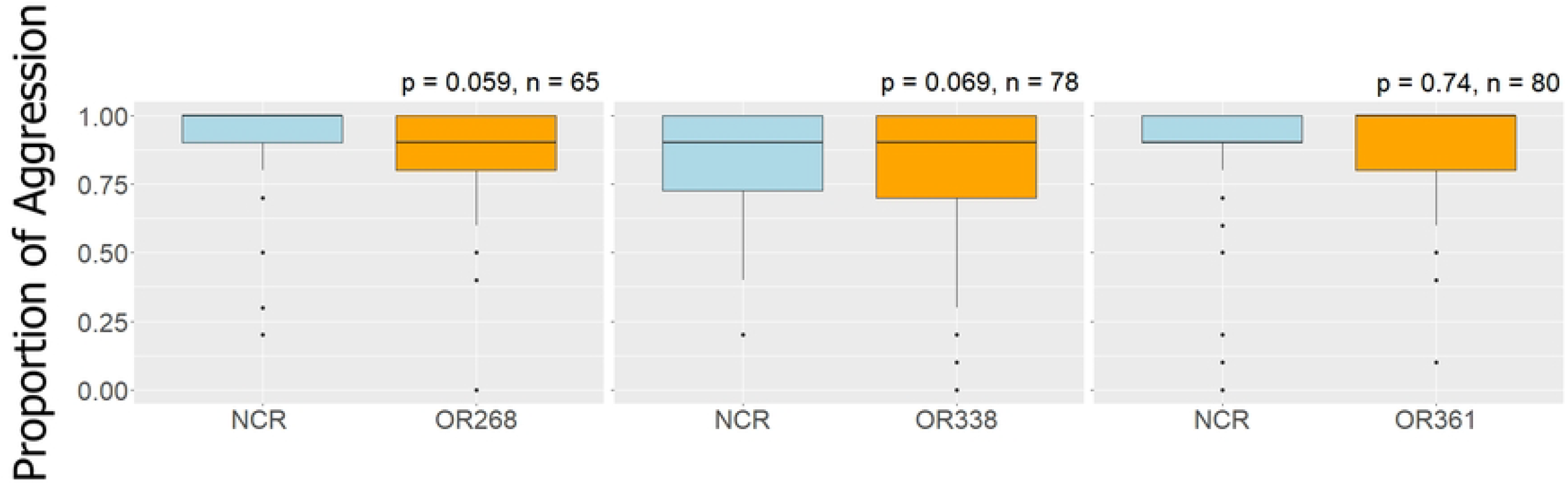
Aggression assay boxplots of 1 µg/µL OR dsRNA vs. NCR dsRNA. ORs *LhOr268* and *LhOr338* show potential mild loss in aggression in OR treated trials, while OR *LhOr361* shows no difference in aggression. Efficacy of tuning OR knockdown on aggression is uncertain and observed changes in behavior tend to be very small.

**Figure 4.**
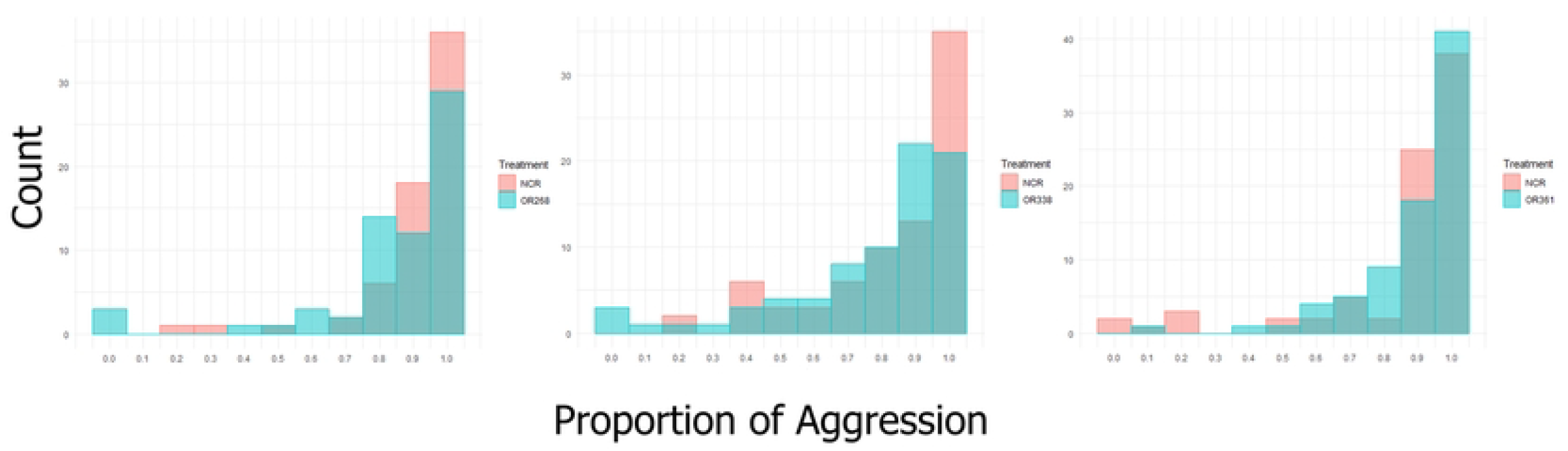
Aggression assay histograms of 1 µg/µL OR dsRNA vs. NCR dsRNA. ORs *LhOr268* and *LhOr338* show similar pattern shift in aggression reduction as displayed in *orco*-treated ants. OR *LhOr361* showed no meaningful shift in patterns of aggression between ants treated with *LhOr*361 dsRNA and NCR dsRNA.

## 5 Discussion

Results demonstrate that dsRNA can cross the midgut tissue and inhibit gene expression even in genes that only show expression in tissues at the body periphery, as exhibited by dsRNA yielding a reduction in *orco* gene expression in antennal tissues. This reduction produced a reduction in aggressive behavior, which provides similar results to the total loss of aggression when treated with *orco* antagonists (Ferguson et al., 2020). Targeting tuning ORs might produce a mild decrease in aggression in some tuning ORs, but evidence remains uncertain. Given the size of the nine-exon OR clades thought to be involved in nestmate recognition, a mild reduction of aggression in response to loss in normal function of a single tuning OR would be expected, potentially generating an effect too small to measure with aggression assays. However, qPCR showed no detectable knockdown of expression of these genes. There are two potential explanations for this lack of knockdown. First, evidence thus far indicates that tuning ORs are expressed within a small number of sensilla (Clyne et al., 1999). Given that much of the dsRNA seems to stay in the midgut, there might be insufficient transfer into the hemolymph to saturate all antennal tissues and ensure the correct ORNs uptake the dsRNA, in which case more research is needed into dsRNA transfer across the midgut to improve transfer efficiency. The alternative explanation is that the knockdown effect of the dsRNA is being masked. Gene expression work in the clonal raider ant, *Ooceraea biroi*, determined that ORs in tandem arrays are transcribed, but only the first gene is exported to the cytoplasm for expression (Brahma et al., 2023). If OR expression in *L. humile* works similarly, the abundance of OR transcripts in the ORN nucleus may be out of reach of the RISC complex and therefore masking the effects of gene knockdown when conducting qPCR analysis. A more quantitative way to measure the effect of tuning OR knockdown would be to conduct coupled gas chromatography-electroantennography (GC-EAG) on treated antennae compared to control, to look for changes in antennal detection of cuticular hydrocarbon components (D’Ettorre, 2004; Shi et al., 2017). This analysis would also provide a more practical method for characterizing the CHC ligands that bind to a specific odorant receptor. Current methods for investigating OR function generally rely on mutagenesis, either using CRISPR to generate a knockout mutant, or through introducing the gene into *Drosophila* fly lines, *Xenopus* oocytes, or human embryonic kidney (HEK293) cell lines (Nakagawa and Touhara, 2013; Roberts et al., 2021; Slone et al., 2017; Yan et al., 2017). RNA interference represents a much less labor- and time-intensive process for conducting functional characterization of genes and allows for the investigation of gene function in its original context. This is particularly necessary in a eusocial insect like the Argentine ant, where the reproductive division of labor makes cultivating a viable population of mutants incredibly difficult.

Additionally, since learning colony odor takes place upon pupal eclosion, any attempt to identify the influence of an OR on nestmate recognition via mutagenesis is not possible (Sturgis and Gordon, 2011). Knockdown of OR function in the adult stage is necessary to investigate how ORs regulate nestmate recognition.

## References

Brahma, A., Frank, D.D., Pastor, P.D.H., Piekarski, P.K., Wang, W., Luo, J.-D., Carroll, T.S., Kronauer, D.J.C., 2023. Transcriptional and post-transcriptional control of odorant receptor choice in ants. Curr. Biol. CB 33, 5456-5466.e5. 10.1016/j.cub.2023.11.025

Clyne, P.J., Warr, C.G., Freeman, M.R., Lessing, D., Kim, J., Carlson, J.R., 1999. A Novel Family of Divergent Seven-Transmembrane Proteins: Candidate Odorant Receptors in Drosophila. Neuron 22, 327–338. 10.1016/S0896-6273(00)81093-4

D’Ettorre, P., 2004. Does she smell like a queen? Chemoreception of a cuticular hydrocarbon signal in the ant Pachycondyla inversa. J. Exp. Biol. 207, 1085–1091. 10.1242/jeb.00865

Dittmann, M.A., Buczkowski, G., Scharf, M., Harpur, B.A., 2024. Comparative transcriptomics and phylostratigraphy of Argentine ant odorant receptors. PLOS ONE 19, e0307604. 10.1371/journal.pone.0307604

Doudna, J.A., Charpentier, E., 2014. The new frontier of genome engineering with CRISPR-Cas9. Science 346, 1258096. 10.1126/science.1258096

Ferguson, S.T., Park, K.Y., Ruff, A.A., Bakis, I., Zwiebel, L.J., 2020. Odor coding of nestmate recognition in the eusocial ant Camponotus floridanus. J. Exp. Biol. 223, jeb215400. 10.1242/jeb.215400

Liang, D., Silverman, J., 2000. “You are what you eat”: Diet modifies cuticular hydrocarbons and nestmate recognition in the Argentine ant, Linepithema humile. Naturwissenschaften 87, 412– 416. 10.1007/s001140050752

McKenzie, S.K., Fetter-Pruneda, I., Ruta, V., Kronauer, D.J.C., 2016. Transcriptomics and neuroanatomy of the clonal raider ant implicate an expanded clade of odorant receptors in chemical communication. Proc. Natl. Acad. Sci. 113, 14091–14096. 10.1073/pnas.1610800113

Nakagawa, T., Touhara, K., 2013. Functional Assays for Insect Olfactory Receptors in Xenopus Oocytes, in: Touhara, K. (Ed.), Pheromone Signaling: Methods and Protocols, Methods in Molecular Biology. Humana Press, Totowa, NJ, pp. 107–119. 10.1007/978-1-62703-619-1_8

Obin, S., 1986. Nestmate Recognition Cues in Laboratory and Field Colonies of Solenopsis invicta (Buren) (Hymenoptera: Formicidae): Effect of Environment and Role of Cuticular Hydrocarbons. J. Chem. Ecol. 12, 11.

Roberts, R.E., Yuvaraj, J.K., Andersson, M.N., 2021. Codon Optimization of Insect Odorant Receptor Genes May Increase Their Stable Expression for Functional Characterization in HEK293 Cells. Front. Cell. Neurosci. 15.

Roulston, T.H., Buczkowski, G., Silverman, J., 2003. Nestmate discrimination in ants: effect of bioassay on aggressive behavior. Insectes Sociaux 50, 151–159. 10.1007/s00040-003-0624-1

Shi, Q., Lu, L., Lei, Y., He, Y., Chen, J., 2017. Gland Origin and Electroantennogram Activity of Volatile Compounds in Ghost Ants, Tapinoma melanocephalum (Hymenoptera: Formicidae) and Behavioral Response to (Z)-9-Nonadecene. Environ. Entomol. 46, 1374–1380. 10.1093/ee/nvx164

Slone, J.D., Pask, G.M., Ferguson, S.T., Millar, J.G., Berger, S.L., Reinberg, D., Liebig, J., Ray, A., Zwiebel, L.J., 2017. Functional characterization of odorant receptors in the ponerine ant, Harpegnathos saltator. Proc. Natl. Acad. Sci. 114, 8586–8591. 10.1073/pnas.1704647114

Smith, A.A., Liebig, J., 2017. The evolution of cuticular fertility signals in eusocial insects. Curr. Opin. Insect Sci., Vectors and medical and veterinary entomology * Social insects 22, 79–84. 10.1016/j.cois.2017.05.017

Smith, C.D., Zimin, A., Holt, C., Abouheif, E., Benton, R., Cash, E., Croset, V., Currie, C.R., Elhaik, E., Elsik, C.G., Fave, M.-J., Fernandes, V., Gadau, J., Gibson, J.D., Graur, D., Grubbs, K.J., Hagen, D.E., Helmkampf, M., Holley, J.-A., Hu, H., Viniegra, A.S.I., Johnson, B.R., Johnson, R.M., Khila, A., Kim, J.W., Laird, J., Mathis, K.A., Moeller, J.A., Muñoz-Torres, M.C., Murphy, M.C., Nakamura, R., Nigam, S., Overson, R.P., Placek, J.E., Rajakumar, R., Reese, J.T., Robertson, H.M., Smith, C.R., Suarez, A.V., Suen, G., Suhr, E.L., Tao, S., Torres, C.W., van Wilgenburg, E., Viljakainen, L., Walden, K.K.O., Wild, A.L., Yandell, M., Yorke, J.A., Tsutsui, N.D., 2011. Draft genome of the globally widespread and invasive Argentine ant (Linepithema humile). Proc. Natl. Acad. Sci. U. S. A. 108, 5673–5678. 10.1073/pnas.1008617108

Sturgis, S.J., Gordon, D.M., 2011. Nestmate recognition in ants (Hymenoptera: Formicidae): a review. Myrmecol. News 10.

Suh, E., Bohbot, J.D., Zwiebel, L.J., 2014. Peripheral olfactory signaling in insects. Curr. Opin. Insect Sci., Pests and resistance/Parasites/Parasitoids/Biological control/Neurosciences 6, 86–92. 10.1016/j.cois.2014.10.006

Trible, W., Olivos-Cisneros, L., McKenzie, S.K., Saragosti, J., Chang, N.-C., Matthews, B.J., Oxley, P.R., Kronauer, D.J.C., 2017. orco Mutagenesis Causes Loss of Antennal Lobe Glomeruli and Impaired Social Behavior in Ants. Cell 170, 727-735.e10. 10.1016/j.cell.2017.07.001

Wilson, R.C., Doudna, J.A., 2013. Molecular Mechanisms of RNA Interference. Annu. Rev. Biophys. 42, 217–239. 10.1146/annurev-biophys-083012-130404

Yan, H., Liebig, J., 2021. Genetic basis of chemical communication in eusocial insects. Genes Dev. 35, 470–482. 10.1101/gad.346965.120

Yan, H., Opachaloemphan, C., Mancini, G., Yang, H., Gallitto, M., Mlejnek, J., Leibholz, A., Haight, K., Ghaninia, M., Huo, L., Perry, M., Slone, J., Zhou, X., Traficante, M., Penick, C.A., Dolezal, K., Gokhale, K., Stevens, K., Fetter-Pruneda, I., Bonasio, R., Zwiebel, L.J., Berger, S.L., Liebig, J., Reinberg, D., Desplan, C., 2017. An Engineered orco Mutation Produces Aberrant Social Behavior and Defective Neural Development in Ants. Cell 170, 736-747.e9. 10.1016/j.cell.2017.06.051

